# Normalizing the brain connectome for communication through synchronization

**DOI:** 10.1101/2020.12.02.408518

**Authors:** S. Petkoski, V.K. Jirsa

**Affiliations:** Aix-Marseille Univ, Inserm, INS, Institut de Neurosciences des Systèmes, 13005 Marseille, France

**Keywords:** Synchronization, Time-delays, Normalized connectome, Spectral activation patterns, Oscillations, Resting state networks

## Abstract

Networks in neuroscience determine how brain function unfolds, and their perturbations lead to psychiatric disorders and brain disease. Brain networks are characterized by their connectomes, which comprise the totality of all connections, and are commonly described by graph theory. This approach is deeply rooted in a particle view of information processing, based on the quantification of informational bits such as firing rates. Oscillations and brain rhythms demand, however, a wave perspective of information processing based on synchronization. We extend traditional graph theory to a dual particle-wave-perspective, integrate time delays due to finite transmission speeds and derive a normalization of the connectome. When applied to the data base of the Human Connectome project, we explain the emergence of frequency-specific network cores including the visual and default mode networks. These findings are robust across human subjects (N=100) and are a fundamental network property within the wave picture. The normalized connectome comprises the particle view in the limit of infinite transmission speeds and opens the applicability of graph theory to a wide range of novel network phenomena, including physiological and pathological brain rhythms. These two perspectives are orthogonal, but not incommensurable, when understood within the novel here proposed generalized framework of structural connectivity.

**AUTHOR SUMMARY:** All networks are composed of nodes and links, forming the structural frame, in which communication occurs. We demonstrate that graph theoretical tools make the implicit assumption of information transmission via exchange of bits, suggesting that the stronger connected nodes are more impactful upon the remainder of the network. This corollary does not extend to communication through oscillations, which is the prominent information carrier in brain networks. We extend traditional network analysis to the oscillatory domain and derive a novel network normalization including descriptive metrics. Along the prototypical example of the brain as a network, we illustrate the consequences of this novel approach and demonstrate that the normalization robustly explains the emergence of the prominent frequency-specific network cores, which cannot be understood within the traditional framework.

## INTRODUCTION

Network theory significantly advanced our understanding of complex processes in nature, ranging from gene regulation of protein interactions (*1*), coordination of brain activity (*2*) to social networks (*3*). Connectivity is the dominant concept that shapes the capacity of a network to transmit information (*4*) and is described by its topological and statistical properties (*5*). In neuroscience, network theory has been applied on several scales, including microscopic neocortical microcircuitry (*6*), macroscopic structural connectivity – connectome (*7*), and networks of coordinated brain activity, also called functional connectivity (*8*). Models of brain networks based on connectomes (*9*) have demonstrated individual predictive power in resting state paradigms (*10*), cognitive tasks (*11*) and brain disease such as epilepsy (*12*). In all these applications, there remains a deeply rooted understanding of signal transmission in which bits of information are transmitted between network nodes, independently of the underlying oscillations and brain rhythms. But, as the particle picture is complemented by a wave picture in physics, similarly in neuroscience there are oscillatory processes deeply implicated in healthy and pathological brain activation (*13*), such as cognitive functions (*14*) or aberrant discharges in epilepsy (*12*). How such rhythms support the communication through the phenomena such as phase and frequency dependance does not have a fundamental description within the particle view of information processing, which describes network processes by activations, but does so in the wave picture that imposes a language in terms of synchronization.

The importance of the particle-wave duality in network neuroscience becomes evident when considering signal transmission delays, present in large scale brain networks and ranging on the order of 10-200 ms (*15*). As physiological rhythms in the brain lie in the same range (*16*), shifts in arrival times due to delays have minimal to no effects in the particle view, but result in frequency dependent effects in the wave view, where a synchronized pair of oscillators may switch from full synchrony to anti-synchrony. As such, frequency and time delays become inseparable properties of the network and together with the connectome determine the synchronization and nodes’ activity (*17, 18*). This also confirms previous computational studies about the effects of the space-time structure of the brain on its emergent dynamics (*19*), and could also determine the spectrally dependent information processing capacities based on synchrony (*20*).

Current state of the art network neuroscience is mainly focused on the topological aspects in describing the connectivity of the brain (*2, 21*), thus limits itself to the particle view only and omits space and physical distances between the interacting units (*22*). Possible exceptions from this are the cases when structural network motifs are shown to be related with the functional hubs due to the time-delays (*23*), but without spectral specificity. In the rare cases when lengths of the links are studied (*24*), it is still done in static manner, without consideration for the impact of the delays on the emerging dynamics, thus critically demanding the extension of network neuroscience to the wave picture. The only attempts to study negative links between brain regions were limited to the clustering of functional data with negative correlations (*22*), without any application to the structural links, for which there was never a mechanistic explanation that would render them effectively inhibitory for the network synchronization.

## RESULTS

### Particle versus wave representation for network dynamics

The particle-wave dichotomy for network communication appears in networks dynamics due to the finite transmission velocities for the signals between spatially distributed nodes. The governing equation reads

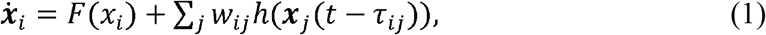

where ***x***_*i*_ is a vector of states, *F* is a non-linear function, and *h* is the coupling function. Within the particle view, ***x*** is a one-dimensional scalar variable such as firing rate, rendering the interactions to be frequency-independent, Fig. 1 (A). Consequently, the strongest flow of information and the activity is generally along the links with the strongest coupling weight (*25*), although there are cases when hubs are significantly more influential (*26, 27*). In addition, time-delays are less important during single events that occur independently of the possible underlying oscillatory activity (middle plot of Fig. 1(A)), such as the action potentials that make up subsequent metrics such as firing rate. Similar examples of particle-like interactions happen during strong perturbations and stimulations (*28*), or when the coupling is no longer weak and bifurcations occur, such as during seizure propagation in epilepsy (*29*).

**Fig. 1.**
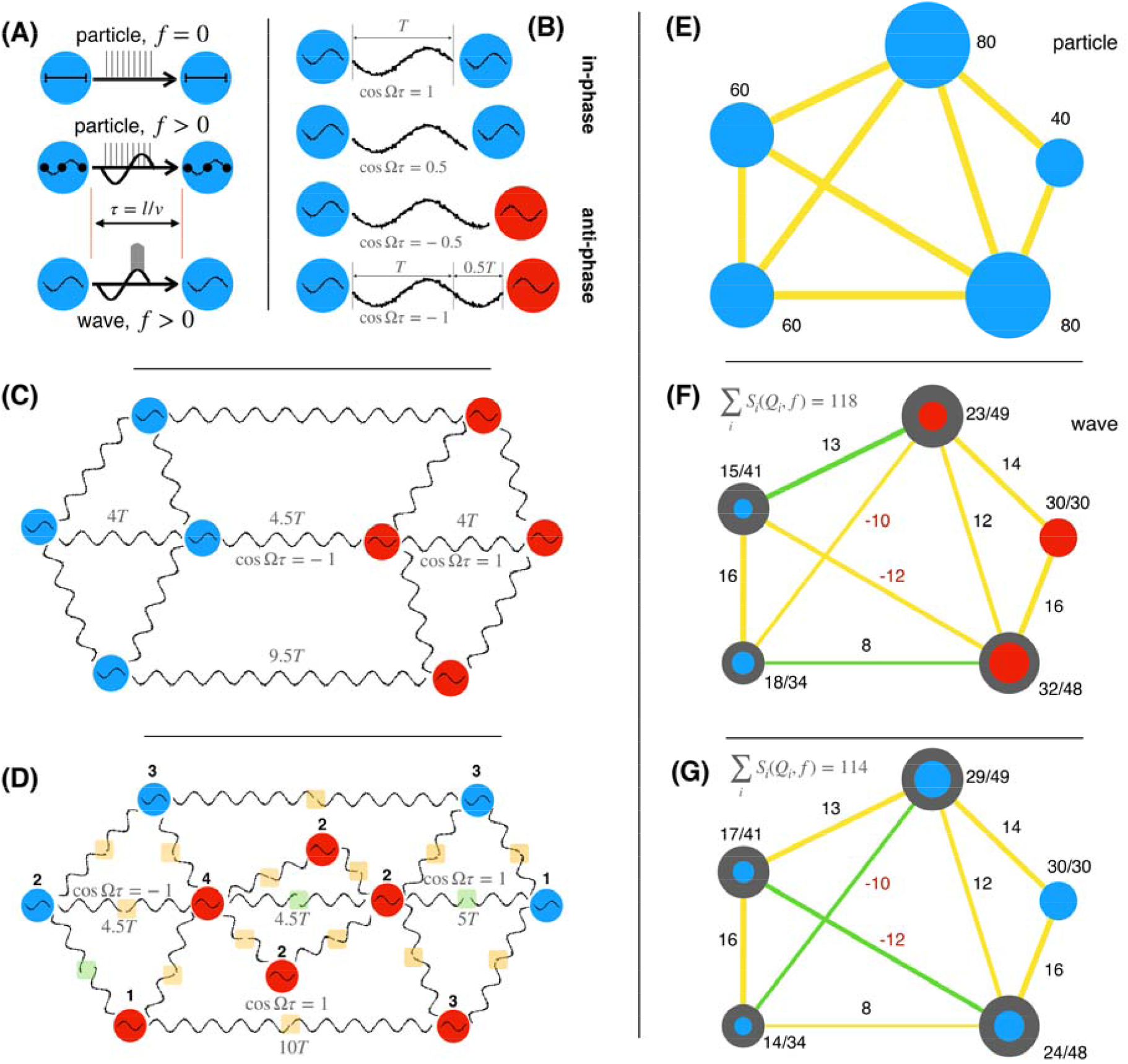
Spatio-temporal network organization and wave coupling weights for synchronized networks. (A) Particle (static or frequency-independent) and wave (synchronization-dependent) interactions with delay *τ*, due to distance *l*, and propagation velocity *v*. (top) Transmission of packets over non-oscillatory local dynamics; (middle) a particle type of transmission that does not depend on the coherence (e.g. during strong perturbation) and is thus independent of the underlying oscillatory dynamics; (bottom) the communication is dependent on the synchronization. (B-D) In-and anti-phase synchronization for fixed frequency and different time-delays for 2 oscillators (B), and for networks with a resonant (C) and non-resonant (D) spatio-temporal alignment of the oscillators. Nodes with same color are in-phase with each other, and anti-phase with others, as also illustrated with the waveform. Time-delays are shown relatively to the period of oscillations *T*, and the normalization factor is shown for some of the links. (D) Midpoints of links with in-phase oscillations are highlighted yellow, and green if anti-phase, and the sum of all positively and negatively contributing links (yellow minus green links) to each node is also shown. (E-G) Wave coupling weights for a particle (or static) case (E) and for an optimal, (F) and suboptimal (G) phase arrangements of nodes. Links contributing positively to the synchronization are yellow, and green are negative contributions. Lines have width proportional to 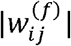 (shown next to the links) and the size of the circles corresponds to the nodes spectral capacity (dark) and strength (colored), both shown for each node.

Networks of oscillators, on the other hand, are often used to study dynamical systems for which the local activity is multidimensional and nonlinear (*30*). They have been conceptualized to be responsible for the communication in the brain through coherence (*20*) and synchronization (*31*), Fig. 1 (A), but this aspect is still unlinked on the network level to the features of the structural links, i.e. weights and time delays, that shape the synchronization of the brain network (*18*). For weak couplings, dynamics of the nonlinear oscillatory system in Eq. (1) are captured by phase models, of which the simplest and the most elaborate due to the analytical tractability is the Kuramoto model (*32*). For synchronization at frequency Ω, the model reads (*33*)

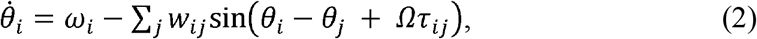

where *θ*_*i*_ denotes the phase of the *i-*th node oscillating with a natural frequency *ω*_*i*_. Note that if time-delays were short so that they would not appear explicitly in the phase model (*33*), then Eq. (2) would have still represented the phase model for the case of synchronization.

From Eq. (2) follows that when delays are negligible compared to the time-scale of the system, the interactions are governed only by the weights, as in the particle case. Due to better tractability and the readily applicable tools from the graph theory, this has been the default network representation in neuroscience. However, when the time-delays are comparable to the time-scale of the intrinsic oscillations, they need to be included in the analysis of network dynamics. This system can only be solved numerically, in order to unveil the activation patterns at different frequencies as there is no alternative graph theoretical approach that integrates the topology and the impact of the time-delays on the oscillations. Hence the systematic effect of the time-delays on the emergent dynamics in Eq. (1) is concealed, beyond the synchronizabillity and the order parameter which can be calculated using the Ott and Antonsen ansatz (*34*), but only for all to all couplings and spatially decomposed delays (*35*).

To go beyond the static representation of networks, we use the insight that the impact of the direct link in the phase difference between oscillators can be separated from the rest of the network (see the Model section), leading to solutions given as

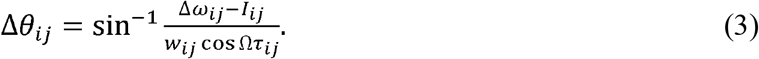

Here Δ*ω*_*ij*_ = (*ω*_*i*_ − *ω*_*j*_)/2, and normalized wave coupling 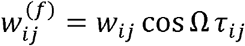 describes the impact of the direct link between the oscillators. The influence of all the other links of these two nodes is contained in *I*_*ij*_. The cosine term vanishes for particle-like communication Ω*τ*_*ij*_ → 0, simplifying Eq. (3) to the case of no delays, when weights are the only factor shaping the network dynamics. For interactions of synchronized oscillations, wave coupling 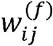 unifies spatio-temporal aspects of each link by modulating weights with the frequency-specific impact of the time-delays. This reflects duality in the network interactions, which in the wave case is reliant on the timings of the waveforms arriving from the distant nodes, Fig. 1 (B-D).

As illustrated in Fig. 1 (B), two delay-coupled oscillators can synchronize either in-phase or anti-phase (*17, 30*), depending on the sign of cosΩ*τ*. The former is stable for positive wave couplings, the latter for negative. The same is true for networks (*17*) and not limited to phase oscillators (*18*). Time-delays can be perfectly spatially distributed to cause maximum synchronization at a given frequency Fig. 1 (C), but a more realistic scenario is when their distribution causes some links to decrease the network synchronization, i.e. if 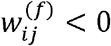 for in-phase nodes, or if 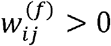 for anti-phase. One possibility here is the oscillators rearranging their phases to minimize this disturbance Fig. 1 (D). From here, to identify the skeleton of the wave synergy we define a spectral strength and capacity for each network node

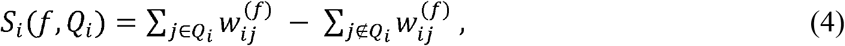

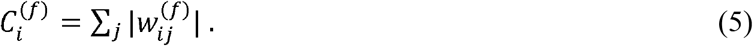

In the static case, both metrics yield the topological node strength, Fig. 1 (E). *S*_*i*_(*f, Q*_*i*_) is adapted from the particle case node strength (*22*) by accounting for possible inhibitory contribution of some links to the overall synchronization, i.e. negative wave coupling from an in-phase node, or a positive one from an anti-phase node, Fig. 1 (D, F, G). The cluster *Q*_*i*_ thus contains all the nodes that are in phase with the node *i*. Related to this, 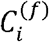 gives the upper bound of node’s synchronizabillity and hence of its *S*_*i*_(*f, Q*_*i*_), Fig. 1 (F, G). Therefore, it shows the strength of the node when all its links contribute positively to its dynamics. As shown in the same plots, different phase arrangements of the nodes change their spectral strength. A hub node can become more peripheral, or vice-versa, while still being constrained by its spectral capacity, which does not change as different arrangements are realized for a fixed frequency.

For simplicity, in the rest of the manuscript we will omit the explicit dependence of the spectral strength on the node-specific cluster *Q*_*i*_, but instead we will refer to a specific in/anti phase arrangements on the network level that imply specific clustering for the nodes. For example, in most of the results we will show results for 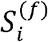 from a clustering that maximizes its sum over the whole network.

### Normalized Connectome and Spectral Dependent Activity of Brain Regions

The wave coupling weights are frequency dependent, and in the case of the connectome this causes activations of different network cores. We assume homogeneous propagation velocity, which is often used as a first approximation, resulting in time delays being defined by the lengths of the links (*9, 17*–*19*), i.e.*τ*_*ij*_ *=l*_*ij*_ /*v*. The cosine term modulates the strength of the links, which can even change the sign, reverting once positive interactions depending on the frequency. This is demonstrated for connectomes of a human (*36*) and mice (*37*), Fig. 2 (A).

**Fig. 2.**
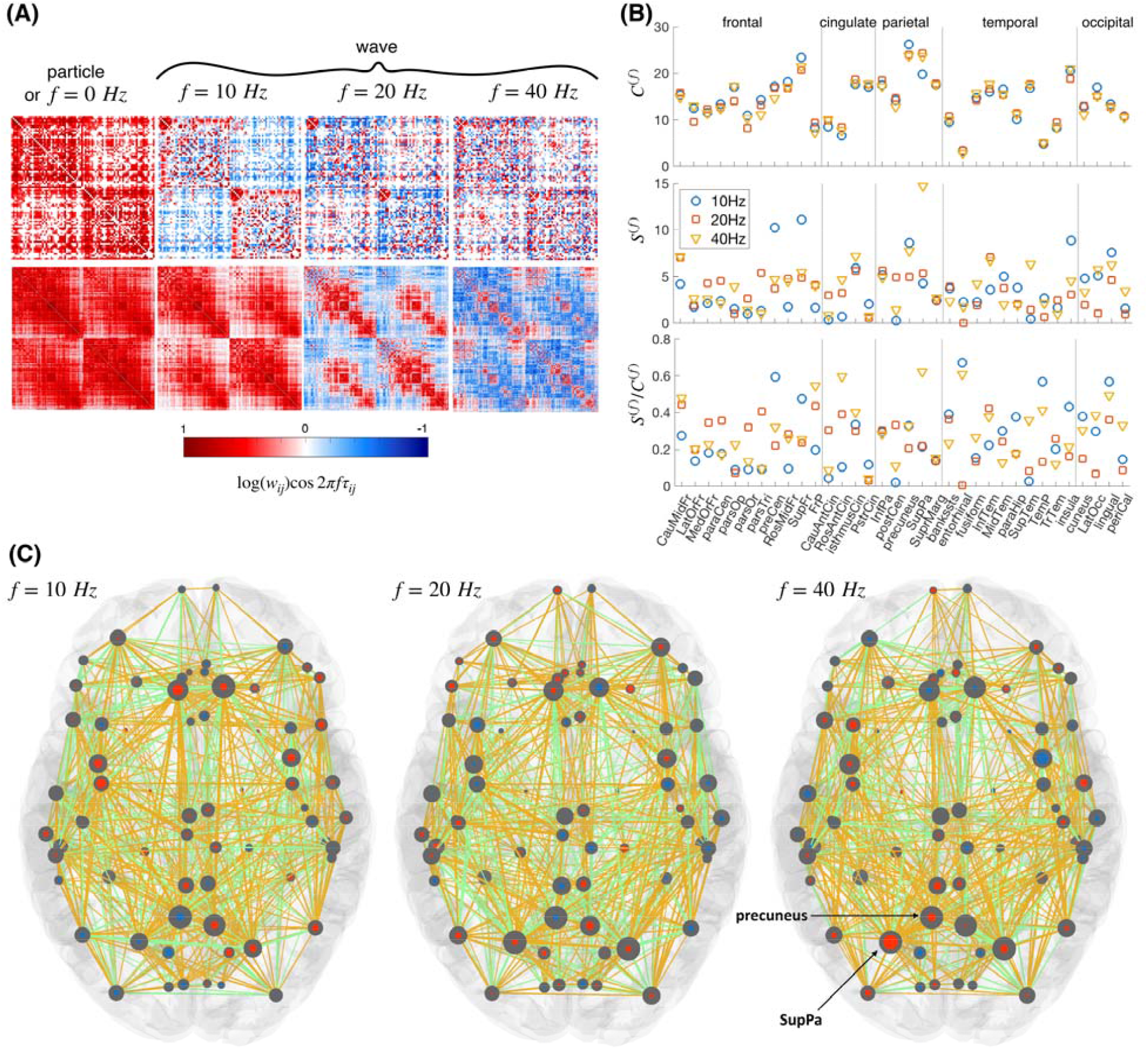
Normalized connectomes at different frequencies. (A) Wave couplings for a human (top) and averaged mice (bottom) connectomes. (B) Spectral strength and capacity, and their ratio for regions of the left hemisphere ordered by lobes for 3 different frequencies. Full names of the regions are shown in Tab. S14. (C) Spectral strength (blue or red, indicating anti-phase arrangements) and capacity (grey) for cortical regions of a human brain. The links with positive contribution to the optimal phase arrangement are yellow, and others are green (c.f. Fig. 1 (B)).

The spectral strength and capacity are illustrated in more details in Fig. 2 (B, C) for three different frequencies, and they are shown for all the brain regions ordered by lobes in Fig. 3. The phase arrangement of the nodes is obtained by maximizing the overall spectral strength of the network (see Methods for more details). For example, the two spectral graph theoretical metrics in Fig. 2 (B, C) illustrate that the highest spectral capacity for regions of cingulate are at 20 Hz and their strength is lowest at 10 Hz, while the occipital lobe has highest spectral strength and capacity at 10 Hz. Large spectral capacity implies possibility for a given node to get stronger strength during different phase arrangements, since as illustrated in Fig. 1 (B), even for a network of several nodes it can already become impossible not to have negatively contributing links that inhibit the synchronization. The ratio 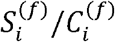 shows what proportion of the weighted links are contributing positively to the spectral strength of the node for the optimal phase arrangement. Hence, the superior parietal (SupPa) region is by far the strongest at 40 Hz due to the more positively contributing links, as compared to the precuneus, which is much weaker at this arrangement, besides having a similar capacity.

**Fig. 3.**
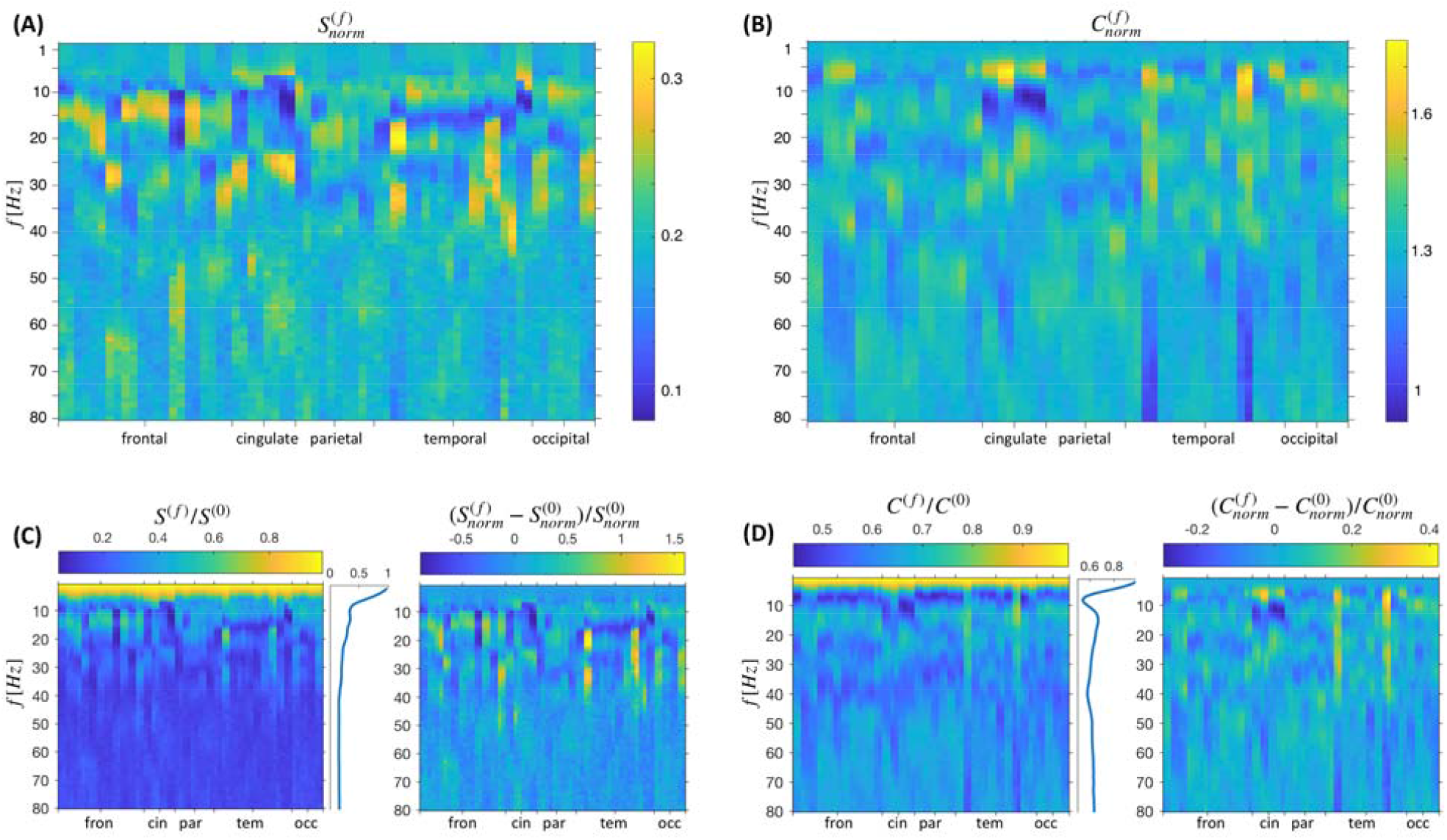
Spectral strength and capacity for brain regions. Mean values are shown over 100 subjects and for frequencies in a step of 1Hz. Brain regions are ordered by lobe with exact ordering within lobes given in Tab. S.14, each right hemisphere region is placed right after its left-hemisphere counterpart. (A) *S*^(*f*)^ and *C*^(*f*)^ normalized across each frequency, (B, C) their ratio and difference compared to the case with no delays, where the former is for the non-normalized metrics, to reveal the frequency dependent mean activity.

At each frequency only the links with spectral strengths above 0.33 of the mean strength are shown for simplicity.

In Fig. 3 relative activation patterns of all the brain regions are shown for the frequencies up to 80 Hz. They are more heterogeneous for the frequencies between alpha and lower gamma, and homotopic brain regions generally show similar activation patterns, due to the high symmetry of the weights and the delays of the connectome. Frequency-normalized values of the spectral strength and capacity unveil the regions with relatively strongest activity at each frequency, whereas the relative change of the activation compared with the particle case is also shown (right plots in Fig. 3 (C, D)). The strength of each region’s activity relative to the particle case, as well as the mean relative activity for the whole brain are also shown in left plots in Fig. 3 (C, D). For every region the strength generally decreases with frequencies, as it is clearly seen in their mean, while the capacity has much steeper decline up to alpha frequencies, when it slightly recovers and stays largely constant.

### Spectral Dependent activity of RSNs and brain lobes

Studies of functional connectivity have demonstrated that signals in brain regions that are related in different tasks, can also correlate even in the absence of an apparent task, hence the notion of resting state networks (RSNs). The most consistent RSNs are default mode network (DMN), visual, sensory/motor (SensMot), auditory, executive control (ExecCont) and frontal-parietal (FronPar) (*38*). In Fig. 4 we show the average spectral strength and capacity of brain regions of 100 subjects, further projected on RSNs and anatomical lobes. Regions with the strongest activity averaged over the subjects are shown in Fig. 4 (A), through the mean 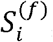, which is normalized at different frequency bands. The same panel contains the boxplots of the strengths of the RSNs of different subjects. The panel (B) of Fig. 4 contains the same metric, but the in-/anti-phase arrangements of the nodes are taken into account through the sign of 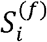, which is now plotted in red-white-blue color code with red regions being anti-phase with the blue. In Fig. 4 (C) the strength at each RSN is projected across the bands, complementing the information about the spectral affinity of RSN. For example, Visual network is by far the most active during alpha frequencies, Fig. 4 (C), even though the alpha frequency also contains a strong activity of the Frontal Parietal network, Fig. 4 (A). Fig. 4 (D) shows the mean and the standard deviation over the 100 subjects of the spectral strength and capacity of each lobe normalized at each frequency. It shows that the robustness over the subjects is much stronger for lower frequency bands (up to around 30 Hz), similarly with the most robust empirically observed spectral activation patterns (*39*–*42*).

**Fig. 4.**
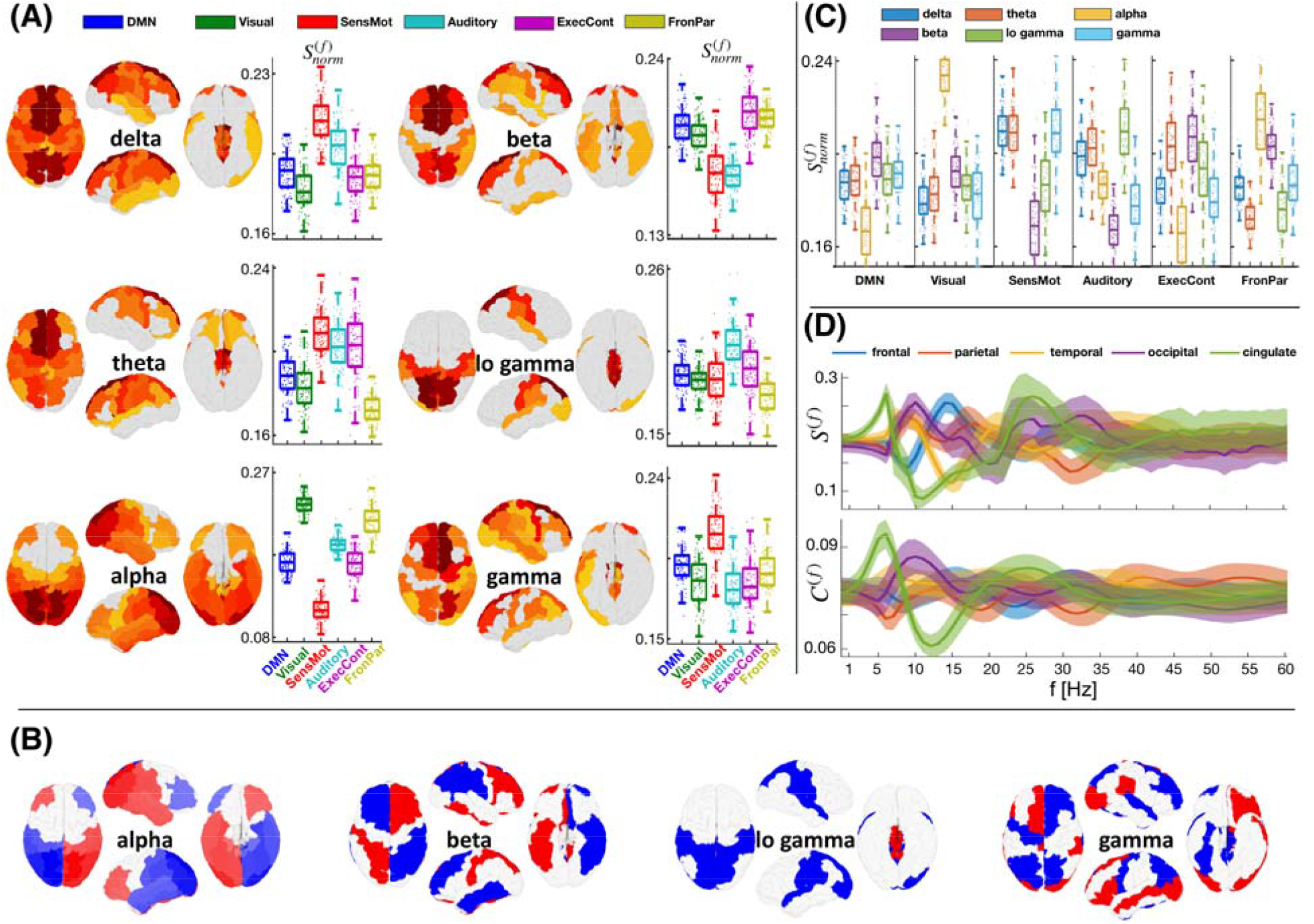
Cortical activation patterns of 100 subjects for different frequencies. (A) Mean spectral strength is shown for different cortical regions (higher for warmer colors) and frequency bands (Tab. S13), and activation of RSNs is projected in the frequencies. (B) In-/anti-phase arrangements of mean 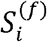 are indicated with red and blue. (C) Relative contribution of the frequency bands per RSN. Boxplots contain the scatter plots of all 100 subjects. (D) Mean spectral strength and capacity (lines) and regions of two standard deviations (shaded) calculated in step of 1Hz for each lobe. The values per given RSN and lobe are normalized by the number of regions in that subnetwork. Brain regions of different RSNs are given in Tab. S.15, and EEG frequency bands are given in Tab. S.13.

Some of the most remarkable and statistically significant (see Tab. S1-S12) activation patterns predicted by the spectral graph theoretical metrics in Fig. 4 are in agreement with the strongest empirically observed phenomena. It is the frequency dependent normalized links that support the visual network areas, which are mainly located in the occipital lobe, to be the most active at alpha frequencies, as it is observed during rest with closed eyes and absence of visual inputs (*43, 44*). Similarly, while activity is dominant in the delta/theta frequencies for the anterior-cognitive state, the posterior-cognitive state is dominated by alpha (*39*). In contrary, the sensory motor and auditory networks are more active at gamma (*45*), whilst DMN, which is mainly in the frontal lobe, is more prominent at beta band (*46*), associated with the idleness of the cognitive and motor setup.

### Numerical confirmation for forward modeling and Linear Stability Analysis (LSA) with Stuart Landau oscillators

The activation patterns predicted by the spectral graph theoretical metrics of the normalized connectome are numerically validated with a linear stability analysis and a forward simulation of brain network model of Stuart-Landau oscillators (*18, 30*) over the same 100 human connectomes, Fig. 5. The model is explored at critical, sub-critical and super-critical regimes, along with noise (see Methods). For consistency, in the simulated results, powers get signs that are based on the in-/anti-phase arrangements of the simulated data, which is obtained from calculating the average phase difference between the regions and checking whether it is larger than *π*. Similarly, for consistent comparison between the metrics, the signs from the phase arrangements either from the data, from the connectome, or from the eigenvalues, are explicitly assigned to the spectral strengths and capacities. Thus, for instance, the spectral strengths respectively become 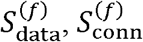 or 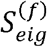.

**Fig. 5.**
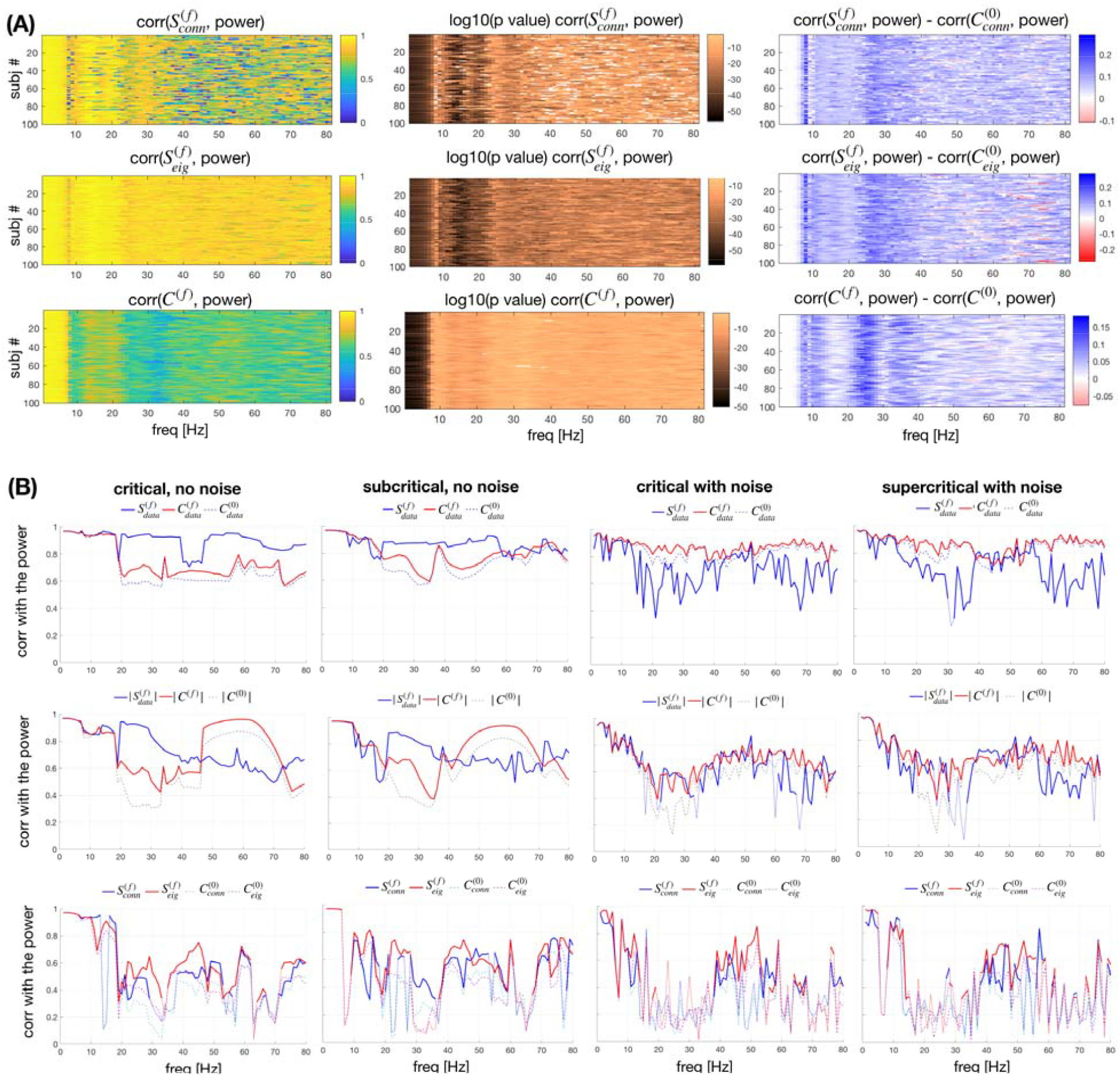
Comparison with forward models based on the normalized and the full connectome. The power across regions from the forward model is compared with the spectral capacity and strength, where the latter is calculated either using the signs of the eigenvectors or using arrangements that maximize its value, and with the null metrics assuming particle-like interactions. The power is calculated (A) using LSA of the brain network model at criticality, constructed over the normalized connectome or (B) with numerical simulations performed for the full connectome, with the brain network model being, sub- (G=17, D=0) and super-critical (G=9, D=1e-5), and at the criticality with noise (G=9, D=1e-5) and without (G=9, D=0). (B) For the full simulations, the most prominent phase arrangement of the simulated data is also used to set the in-/anti-phase arrangements. Correlations of the powers across frequencies (A) for all 100 subjects, as well as the p values, and the difference with the correlations assuming particle-like interactions, and (B) for one subject. Correlations with p<0.01 are transparent in the first 2 columns of (A), and thinner lines for (B).

With the generative model we confirm that these spectral patterns of activity are predicted by the two graph theoretical metrics on the normalized connectome, i.e. the spectral strength and capacity. The former depends on the actual phase arrangement of the oscillators. For this besides the arrangement that maximizes the overall *S*^(*f*)^ (that is based only on the normalized connectome) we also use the signs of the eigenvectors to inform the spectral strength metric about the in/anti phase organization for the specific model. In this dynamical setting the amplitude and the sign of the observed activation pattern for the network coupled through the normalized connectome is guaranteed to be fully captured by the values of the first eigenvector of the LSA (*19, 29*). The signs of the different directions of the eigenvector capture the in-/anti-phase arrangements of the nodes, with the opposite sign regions corresponding to anti-phase synchronized regions.

In Fig. 5 (A), we show the correlation of the leading eigenvectors with spectral strength, calculated using both arrangements, for all 100 subjects. The arrangement implied by the signs of the eigenvectors takes into account the specificity of the model, and hence this case better predicts the activation of a given node. This means that the network is not always arranged in such a manner to yield maximal possible synchronization. The patterns are similar across subjects, and for frequencies larger than 40Hz, there seem to be specific frequencies for different subjects, for which the arrangement of the signs of the eigenvectors is quite different from those maximizing *S*^(*f*)^. Spectral capacity on the other hand, has generally lower predictability than 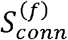, but it is never insignificant or lower than the case without normalization. As shown in the third column of the panel (A), all three spectral metrics exhibit much better correlation than the null metric, which is constructed assuming particle-like interactions at 0Hz. Thus, only the weights of the links are considered, giving the in-strength of the nodes 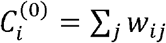. Since the null model cannot yield anti-phase nodes, for more conservative comparison we assign it signs predicted by the arrangements of the nodes with which it is being compared.

In Fig. 5 (B) the predictivity of the spectral metrics is tested by comparing them with the power obtained by simulating the model built with the same oscillators, Eq. (17), but over the full connectome containing both weights and delays, Eq. (18). Moreover, this is extended for the sub-critical and super-critical regimes of the system, along with noise. The comparison shows that each of the wave interactions metrics has consistently higher predictability than its particle-like counterpart across all the frequencies. This holds even for noisy systems far from the criticality, where the correlation is generally larger than 0.5 if the phase arrangements are taken from the data. It is worth noting that for the full system the spectral capacity seems to be more informative than the spectral strength, because the dynamics often becomes non-stationary and multi-stable due to the explicit time-delays (*35*). Nodes might switch from in- to anti-phase and back, and different phase arrangements lead to different spectral strengths over time. As a result, each node’s average relative strength within the network might be better captured by its capacity, than by any given realization of the phase arrangements that has occurred at some point, even if that is the most dominant one. This is especially pronounced when noise is added to the system, making the spectral capacity more informative for the relative activity of the nodes almost for all the frequencies.

### Robustness of the spectral strength

The consistency of the spectral strength of the normalized connectome is tested against different types of uncertainties, which are modeled as rewiring and noise in the connectome, all scaled to have a comparable impact. Boxplots that show the statistics of the leading eigenvector *v* and its absolute value, are shown in Fig. 6 (A) for one frequency and for two different levels of noise. At lower noise, the means of the regional activity shown by the eigenvectors of the null model are still following the original values. The variance of the null models’ activity in this case would sufficiently describe the impact of the noise, and the error that it causes. As the noise increases, the variance of the null model also increases, but in addition for some regions the mean starts significantly differing from the original values. These suggests that to better account the impact of the noise, and the network’s resilience, we need to take into account both the standard deviation *σ*_*i*_ of all realizations of 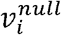, and the error of their mean 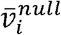 defined as

**Fig. 6.**
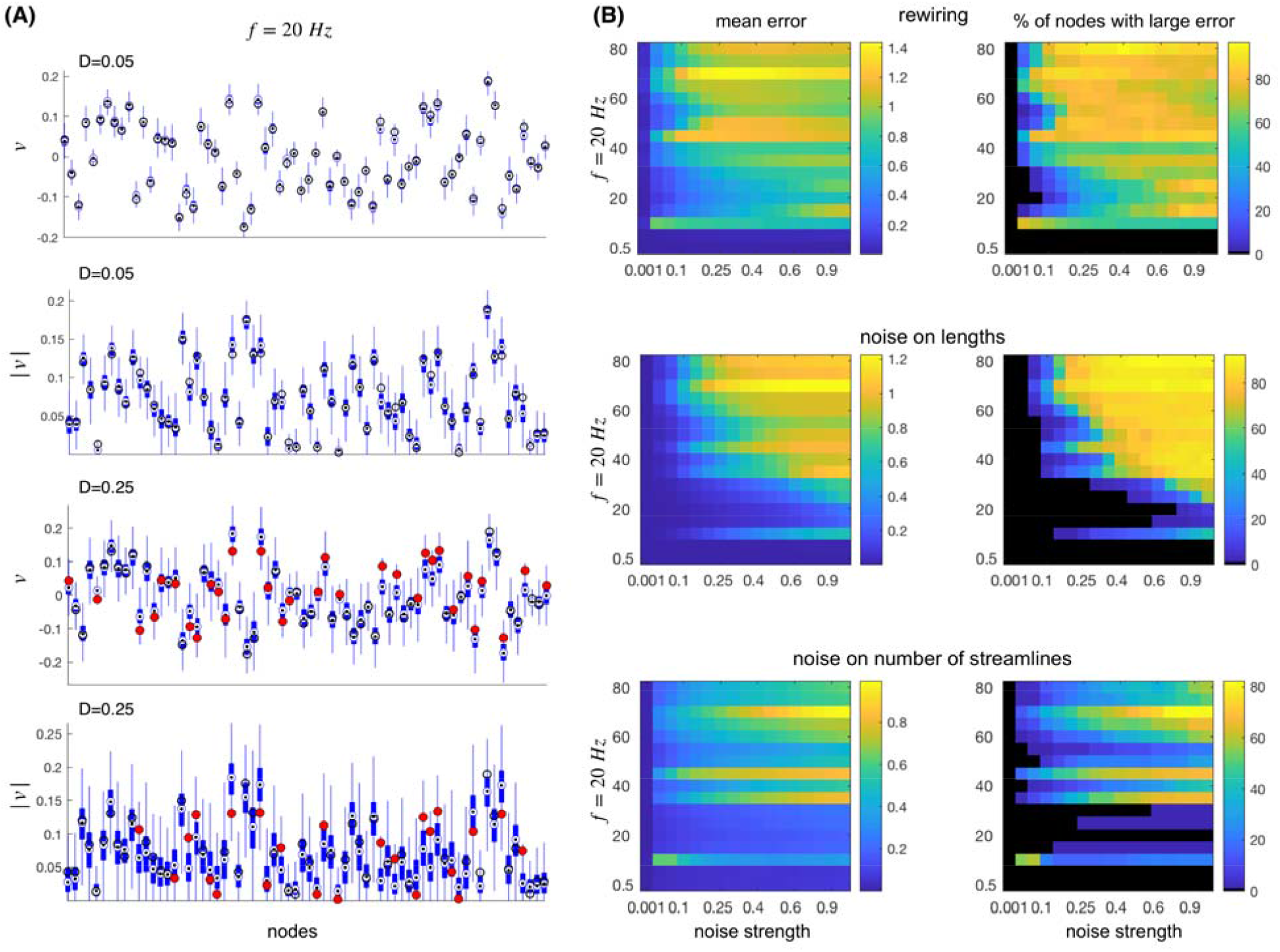
Robustness of the spectral strength. (A) Boxplots showing the statistics of the leading eigenvector and its absolute value for 1000 realizations of 2 different levels of noise added to the lengths at *f* = 20*Hz* on the connectome of the first subject. Black circles are the values for the original connectome, which are filled red for the cases when the error terms Eq. (7), exceed the error threshold. (B) Mean of the errors Eq. (7), over all the regions shown for different noise and frequencies (left column), as well as the ratio of the nodes whose error exceeds the threshold (right column).

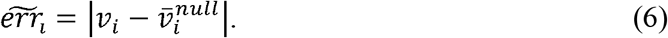

Moreover, the impact of the above error seems to be inversely proportional with the spread of the results for the null model, i.e. smaller the spread, larger the chance for the null model to give an error around its mean. Hence, we arrive to the following definition of the error for each node

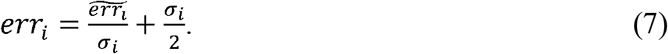

The mean of those errors over the whole network is shown in the first column of Fig. 6 (B). Finally, not all the nodes are equally affected by the noise, and to describe this in the right column plots we depict the proportion of the nodes that have error larger than the threshold set at 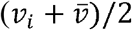. In this way the deviation at each node is evaluated against both, the local, i.e. its own, and the global activity.

Results in Fig. 6 (B) show that the robustness against the noise and rewiring decreases with the frequency, and this is especially evident for the uncertainties of the delays. For frequencies below ∼35 Hz less than half of the nodes have deviations that are considered large for all three types of uncertainties, with rewiring being more sensitive than the noise of the same impact.

## DISCUSSION

Wave metrics offer a new perspective for characterizing dynamic networks. They integrate the wave component of information processing, which was missing in network metrics that were based solely on the connection topology (*2*). The incorporation of signal transmission delays in the connectome’s metrics completes the characterization of the spatiotemporal skeleton, within which oscillatory brain activity can be amplified by the properties of the medium supporting it, i.e. it provides a *corpus resonantiae*. We propose that the activation of certain parts of the brain that are related to different tasks can be explained as being prewired in the anatomy, and we have demonstrated here that the brain connectome has such properties and allows for selective and frequency dependent information processing that could support the differentiation of the brain activity for various processes and frequency bands (*47, 48*). Notably, the wave perspective naturally leads to the formation of overlapping, albeit functionally independent, subnetworks, which cannot be derived within the particle picture. Besides revealing the frequency specific hubness, as another consequence the wave couplings also support the general reduction of activity with increasing the frequencies, which is among the most consistent and well-established features of the brain dynamics (*16*).

The normalization of the connectome that we propose is derived from the expressions for the phase lags between synchronized phase oscillators. These are the only dynamical feature by which functional heterogeneity can arise in this set-up (*17, 49*), due to the structural symmetry breaking of the network interactions (weights and time-delays), or due to the different natural frequencies. Both these aspects are present in Eq. 3, but in this study, we focus on the former. In that respect we show that the in presence of time-delays the weights are effectively scaled (or normalized) by cos Ω*τ*_*ij*_, even though the rest of the network is also needed to fully describe the phase lags and hence the full dynamics. For the case of 2 oscillators, however, the normalized weights are the sole determinant for the synchronized interactions.

Importantly, however, the network models that we introduced are not intended to serve as toy brain models, but to illustrate the meaning of the here derived metrics. We use established models to expose and justify open and hidden assumptions underlying the traditional and extended metrics (the latter being normalized for time delays in the wave perspective). Every metrics relies on an underlying theory/model, even if it is not made explicit. For instance, functional connectivity relies via covariance on second order metrics (*8*) adapted to linear models (fitting an ellipsoid to the corresponding data cloud), but this does not preclude its application to nonlinear network models, in fact in practice this is often the rule. In the same spirit, we motivate the particle and wave metrics by using rate and Kuramoto oscillator equations, without the claim that these actually represent brain network models.

It is worth noting that studying the synchronizabillity of the brain networks with time-delays could complement our framework for the study of spectrally dependent emergent functional heterogeneity, beyond the forward modelling. A possible approach here would be to use recent extensions of the Ott and Antonsen ansatz (*34*) that allow studying noisy networks, either through circular cumulants (*50*), or by using Daido order parameters (*51*). These could be applied on the spatially decomposed time-delays as a first approximation of the spatio-temporal structure of the connectome (*35*). Another direction would be to study the stability of the synchronized states over the network of wave couplings which can be positive and negative (*52*).

During tasks the spectrally dependent activation patterns are also constrained by stimuli that in case of working memory visual tasks for example, cause gamma synchronization in frontal regions (*53*), which is missing from our predictions at rest and in absence of stimuli. The offered framework also do not account for communication across different frequencies, even though it could be further generalized to other types of *n:m* synchronization (*30*). Having data informed different intrinsic dynamics is also a feature that would increase the predictive value of any generative model for the brain dynamics, but this would need to be biased by the data, which is specific of the brain activity (*54*–*56*). This is however still to be generalized in a graph theoretical framework. Similarly, our framework considers the activity at each frequency as fixed (even though they are often time-dependent (*57*–*60*) and intermittent (*61*)) and as a separate process. The latter might lead to omitting certain emergent properties in the cases when the same process has clear multimodal frequency content as often seen in frequency and scale dependent metrics from information theory (*62*–*64*) and time-frequency representation (*65*).

Beyond synchronization, frequency-dependent approaches to information processing offer data driven insight into causality of network interactions (*66*). Applying transfer entropy (*67*) reveals that high degree neurons feed cortical computations by receiving more information from high out-degree neurons (*26*), indicating an asymmetry of information transfer between hubs and low-degree nodes (*27*). Spectral and multiscale Granger causality (*64, 68*) are similarly used to describe the information flow in networks, and they demonstrate the nonlinear effects of the structural weights, which can be seen through signatures of the law of diminishing marginal returns (*69, 70*). It would be interesting to compare the network flow that these metrics reveal for the case of synchronization, with the spectrally dependent network core as defined by the normalized connectome.

The predictive value of the normalized connectome for the functional in-strength is further validated through comparisons with numerical simulations of Stuart Landau model. This model is often used to describe mesoscopic neuronal activity (*18, 55, 71, 72*) due to the fact that it encompasses the working points of population rate models (*73*). Correlating the spatial activation patterns with the predictions from the spectral in-strength and capacity shows that the normalized metrics captures the dynamics of the model better than using only the weights, besides the derivations being made on the Kuramoto model that does not contain amplitude dynamics, and that assumes difference instead of additive coupling. Better performance of the wave coupling is consistent across the frequencies, and holds for different working points, such as sub and supercritical, with or without added noise.

The normalization of the connectome unifies its structural weights and time-delays, and consequently it still inherits the known problems of these two modalities. What the number of streamlines mean for the actual effective coupling between brain regions regardless of the task is still lacking a clear response (*8, 74*). In addition, diffusion tractography is known to under sample the interhemispheric links (*75*), and the topology of the derived connectomes is still dependent on the parcellation (*76*). The lack of personalized estimations for the time-delays of the connectome is still unavoidable problem. A possible way forward here could be using averaged delays derived from stimulation (*77*), combined with the personalized lengths of the tracts, even if these datasets are still incomplete. The lack of directionality in the human connectomes that rely on diffusion tractography is also a confounding factor of those connectomes (*10*), which has led to introduction of concepts such as effective connectivity (*8, 74*), even though for our framework the derivations are simplified by the symmetry in the coupling strengths.

Finally, we have only introduced two metrics for the node activity based on the normalized weights, whereas other graph theoretical metrics for the wave interactions could be more informative. However, even for the spectral capacity and strength, it is still not clear which one would have higher predictive value in describing the activation patterns of the brain at rest. Moreover, their systematic and more thorough comparison against empirical data could provide with the answers about which brain regions operate closer to the limits of their capacity. Similarly, the phase arrangements of the brain do not necessarily need to be such that maximizes the overall information transfer across the whole brain, but maybe instead it gives priority at certain areas, at expense of the others.

Transient synchronization between brain areas coupled through delayed interactions has been proposed as mechanism that could support flexible information routing (*78*), which plays a fundamental role for the cognition. Similarly, the implementation of selective communication dependent on coherence has been generalized to Communication Through Coherence (*20*), as one of the possible mechanisms of communication between neuronal networks (*79*), but with no explanations of the origin of the selection of frequency bands and the spatial organization of the involved neuronal populations. The normalization of the connectome completes this view mechanistically, hypothesizing that the spatio-temporal brain structure itself could be at least partly responsible for the observed patterns. Through the notion of wave-type communication, the offered normalization lends legitimacy to concepts of synchronization and coherence in network science and opens up a many new applications ways beyond brain sciences.

## MODEL & METHODS

### Particle versus wave representation for network dynamics

The particle-wave dichotomy appears in networks dynamics due to the finite transmission velocities for the signals between spatially distributed network nodes. Within the particle view, the general network dynamics is governed by

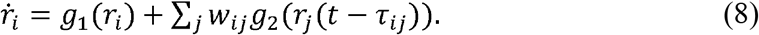

Here *r*_*i*_ is any scalar variable such as the firing rate, *w*_*ij*_ are the weights and *τ*_*ij*_ are the time-delays of the links, and *g*_1_ and *g*_2_ are linear functions for the intrinsic dynamics and coupling respectively. This type of particle-like interactions is frequency-independent and here the information and the activity always flow along the links with the strongest coupling weight.

On the other hand, networks of oscillators are often used to conceptualize and to study dynamical systems for which the local activity is multidimensional and nonlinear (*30*). They have been conceptualized to be responsible for the communication in the brain through coherence (*20*) and synchronization (*31, 78, 80*). In this case the dynamics of the oscillatory network nodes are governed by Eq. (1). When delays are negligible compared to the time-scale of the system, the interactions are still governed only by the weights, as in the particle case. However, when the time-delays are comparable to the time-scale of the intrinsic oscillations, they need to be included in the analysis of network dynamics. As a consequence, this system can only be solved numerically, as there is no alternative graph theoretical approach that integrates the topology and the impact of the time-delays on the oscillations at a given frequency. Hence the systematic effect of the time-delays on the emergent dynamics is concealed and even the full representation of network interactions in Eq. (1), with separated *w*_*ij*_ and *τ*_*ij*_ cannot adequately identify the skeleton of the wave synergy.

For weak couplings, dynamics of the system in Eq. (1) is captured by phase models, which are often used to study the interactions in the brain at different levels (*58, 61*). The simplest and hence most elaborate is the Kuramoto model (*32*). It is widely utilized for describing emergent phenomena in complex systems (*81*), such as the brain (*82*–*84*), with a structure often represented via complex networks (*85*). For synchronization at frequency Ω, the model reads (*33*)

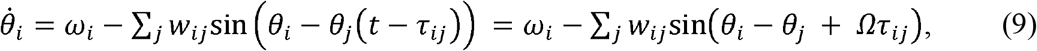

where *θ*_*i*_ denotes the phase of the *i-*th node oscillating with a natural frequency *ω*_*i*_, the dependence on time is implicit, and due to the synchronization *θ*_*j*_ (*t* − *τ*_*ij*_) = *θ*_*j*_ (*t*) + *Ωτ*_*ij*_.

From here, the phase difference between each pair of Kuramoto oscillators is given as

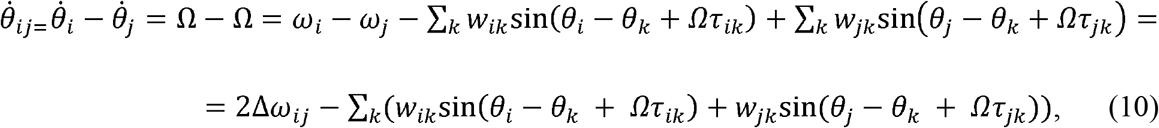

where Δ*ω*_*ij*_ = (*ω*_*i*_ − *ω*_*j*_)/2. Next we introduce Δ*θ*_*ij*_ *= θ*_*i*_−*θ*_*j*_ and we separate the sum, such that

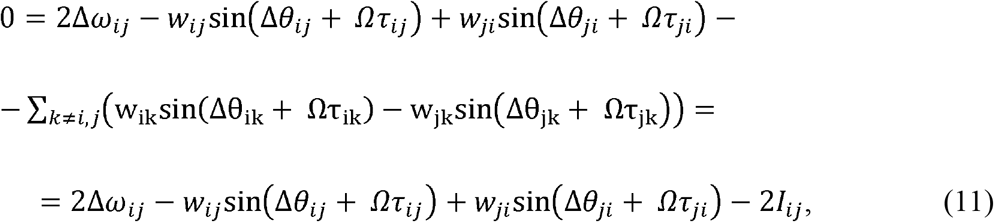

where

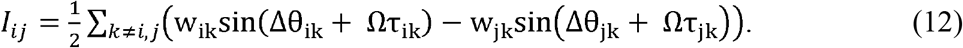

Note that *I*_*ij*_ contains all the other links towards the nodes *i* and *j* apart their direct link.

For symmetric coupling, *w*_*ij*_ *= w*_*ji*_ and *τ*_*ij*_ *= τ*_*ji*_, while Δ*θ*_*ij*_ = Δ*θ*_*ij*_ by definition, hence Eq. (11) becomes

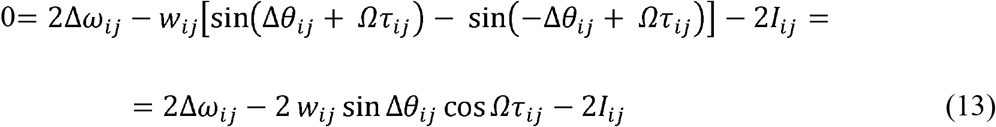

Finally, for the phase difference between the oscillators we get

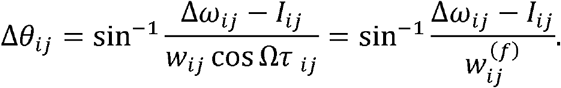

where 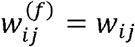 cos Ω*τ*_*ij*_ is the normalized weight containing the impact of the direct link between the pair of oscillators. For particle-like communication in the limit Ω*τ* _*ij*_ → 0 the cosine term and *I*_*ij*_ vanish making the coupling weights the only factor shaping the network dynamics, consistent with the case of no delays. Numerical simulations validate the analytical results for Δ*θ*_*ij*_ by showing that the difference between the analytical and numerical values is small and decreases for smaller integration step, and increase with added noise (Fig. S1).

In the case of asymmetric weights, Eq. (13) would also include a term that accounts for the asymmetry in the interactions *w*_*ij*_ − *w*_*ji*_, and the exact derivation of the expression in Eq. (3) will not be possible anymore. However, if the asymmetry is relatively small, as it is the case for the mice tracer connectome (*37*) for example, then the same expression would approximately hold up to a leading term. As for the human connectomes, they almost exclusively rely on diffusion tractography imaging, which produces links with symmetric weights.

### Limits of the approximation

If we assume normalized connectome, the governing equation of the phase dynamics becomes

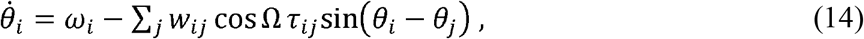

Following the same steps as above, the phase difference between a pair of oscillators reads

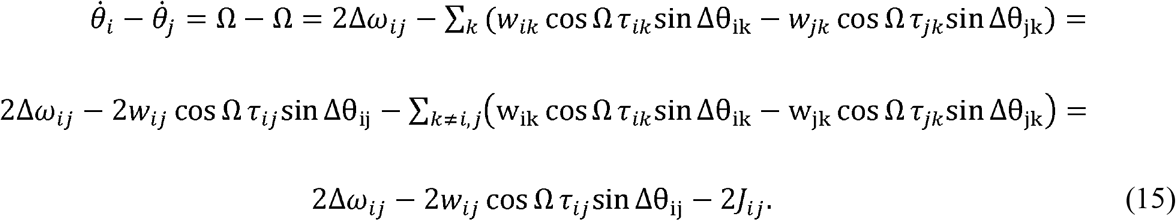

As for the difference between the full model and the approximation using the normalized weights, we get

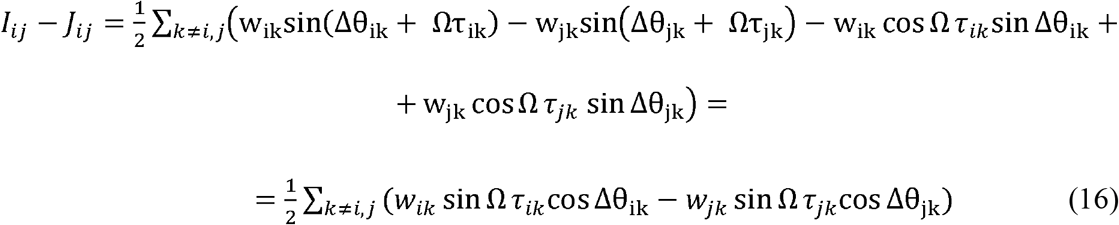

For many symmetric networks, such as the delay-imposed clusters with identical weights (*35*) this difference vanishes.

### Forward modeling and Linear Stability Analysis (LSA)

The normalization of the connectome also allows straightforward application of LSA for the dynamics close to a critical state for a given forward model of the oscillatory activity. To build our brain network model (*17, 18, 29, 55*) over the same 100 human connectomes, we choose Landau Stuart oscillators, as a canonical model for oscillations near Andronov - Hopf bifurcation (*32*), which is also the working point of many neuronal mass models (*86, 87*). The local dynamics from Eq. (1) is then given as

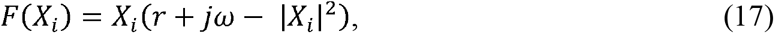

where *X*_*i*_ is a complex number, and the coupling is linear additive with

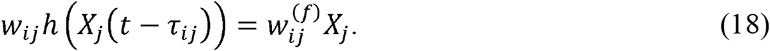

All the parameters, including the frequency *ω* and the distance from the local bifurcation *r* are identical for every oscillator. The stability of this system can be analyzed by following the evolution of infinitesimal perturbations in the tangent space, whose dynamics is ruled by the linearization of Eq. 1 with Eqs. (17 – 18). If the eigenvalues of the Jacobian matrix all have real parts less than zero, then the steady state is stable. For each frequency for which the system is analyzed, we set the parameter *r* to keep the system at the edge of criticality. This is done in an iterative procedure, where *r* is decreased until the system is just above the criticality. In this case there are two identical positive eigenvalues, which contain the same eigenvectors of opposite signs, due to the way how the two dimensions of the local model are coupled.

In addition to the LSA, we carried a numerical simulation over the full connectome of the same model, with the coupling term reading

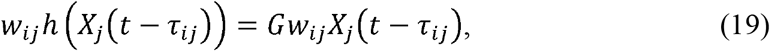

where *G* is a global coupling parameter. Numerical integration is performed using stochastic Heun integration scheme implemented in Matlab, with an integration time-step that is inversely proportional of the natural frequencies and the global coupling, but it is never larger than 0.005s. In some of the analyzed scenarios the dynamics also contains white additive Gaussian, with noise intensity *D*. The critical value of *r* is different for each connectome and each frequency, and sub- and super-criticality are set such that *r* is at 0.2 below or above criticality, respectively.

### Setting the weights for the coupling

Human connectomes were derived from the first release of Diffusion Tractography Imaging of 100 healthy subjects, which are publicly available as part of the Human Connectome project (*36*). The subjects were scanned on a customized 3T scanner at Washington University and the structural connectivity was constructed using a publicly available pipeline (*88*) that applies spherical deconvolution method to a probabilistic streamlines tracking algorithm. The obtained connectome consists of few million tracts spatially averaged to connect 68 cortical regions defined according to Desikan-Kiliany atlas (*89*). For each link, weights are numbers of individual tracts between the regions of that link, and lengths are their averages.

The mouse connectome in Fig. 2 (A) was obtained from the Allen Institute Mouse Brain Connectivity data (*37*), which is integrated in The Virtual Mouse Brain (*90*), and which was already used to model mice brain dynamics (*10, 84*).

The spectral graph theoretical results (Fig. 2-4) and the LSA analysis (Fig. 5 (A) and 6) use the natural logarithms of number of streamlines as weights of the couplings. This is done for better illustration of the findings in order to decrease the heterogeneity between the different nodes. For the full forward model (Fig. 5 (B)), we use the number of streamlines, as it has been more common in the numerical simulations (*17, 18, 29, 55*).

### Phase arrangement of the nodes that maximizes the spectral strength

When the in-/anti-phase arrangement of the nodes is such that *s*^(*f*)^ is maximized, this is referred to as 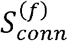, since the arrangement takes into account only the connectome, but not the model. If not specified otherwise in the text, then by the spectral strength *s*^(*f*)^ we refer to 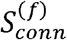. The clusters of in-/anti-phase nodes can be of any size, but the assigning of the clusters is not important (e.g. *i* ∈ *Q*_*i*_ ≡ {1,2, … *N*}\*i* ∈ *Q*_*i*_, and similarly swapping all the red and blue colors in Fig. 1 does not change the results). As a consequence the total number of combinations is 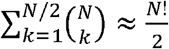 and a brute force procedure cannot be used to identify the clustering that yields the maximum 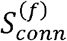. Hence, we developed a heuristic procedure that seems to be able to identify the clusters of maximum spectral strength. In short, it starts with random initial arrangements and then uses three types of cost functions to identify the node that needs to change the cluster. These are:

- The node with smallest *s*^(*f*)^ changes the cluster.
- The node with a smaller *s*^(*f*)^ from the 2 nodes of the link that has the smallest (most negative) internal (from the links within each cluster) 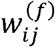.
- The node with a smaller *s*^(*f*)^ from the 2 nodes of the link that has the smallest (most negative) external (from the links between the clusters) 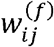.

For better sampling of the different arrangements, a randomness is also systematically added in some of the loops, so that the cost function chooses a suboptimal node or link among some proportion of them that were identified to have the largest cost.

The procedure is run over multiple initializations, until it recovers a cluster that has been already identified. At the end of the runs, the largest spectral capacity that was obtained is set to be 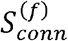. In particular, we start from 10 random initialization for the phases and for each of them the three procedures are repeated one after another until an already visited configuration is achieved, after which the loop is repeated 10 more times with added randomness. Every loop is stopped after arriving to an already visited configuration, and all the visited configurations are then classified according to the highest spectral strength. We have verified that the convergence to the selected configuration occurs even after 10-100 times less samples than we generate with our procedure.

### Projecting the spectral strengths and capacities into RSNs and brain lobes

To better illustrate the patterns of spectral activity we project the spectral strength and capacity in each brain lobe and RSN, Figs. 2-4. For each subject, after calculating them for a given frequency (from 0 to 80 Hz with a step of 1Hz), we calculate the average (per number of regions in the given subnetwork) activity of a given subnetwork (RSN or lobe). In addition, for the results in Fig. 3 and 4 the values of each subnetwork are normalized at each frequency, to focus on the relative activation patterns. The activity of different frequency bands contains the averaged activity of the frequencies within that band (see Tab. S.13). Mean and the standard deviation over the subjects are calculated for each frequency for the activity of the brain lobes.

Note that the averaging over subjects in Fig. 4 (B) is done over the spectral strengths, which explicitly take into account the in-/anti-phase arrangements by assigning them opposite signs. In particular, the regions which are in-phase with regions that have strongest spectral strength over the subjects are set to be always of a positive sign (blue color), as compared with the others which are negative (red). Since during the averaging some nodes can be of opposite phase arrangements over the subjects, the absolute value of the observed activity here, still can be still different from the one shown in Fig. 4 (A).

### Robustness of the spectral strength

To test the robustness of the results, we modified connectomes by either introducing noise or rewiring. We use 3 types of modified connectomes:

- Noise is added to the lengths of the links, with the strength proportional to the mean length of over the links.
- Noise is added to the number of streamlines that are used for calculating the weights. The level of the noise is proportional to the mean number of streamlines of over the links.
- A proportion of the links equivalent to the level of noise is rewired while preserving the distribution of in-strengths (*91*).

For each of this and for each frequency we perform 1000 realizations of the null networks, and then we compute the LSA for each frequency, to obtain the patterns of activity.

## Acknowledgment

This research was supported by the European Union’s Horizon 2020 research and innovation program under grant agreement No. 785907 (SGA2) and 945539 (SGA3) Human Brain Project, and by grant agreement No. 826421 Virtual Brain Cloud.

## Data availability statement

All the data, the codes, and all the relevant information to reproduce the results in the paper will be made available in a public repository.

